# Dynamic and synergistic influences of air temperature and rainfall on general flowering in a Bornean lowland tropical forest

**DOI:** 10.1101/576231

**Authors:** Masayuki Ushio, Yutaka Osada, Tomo’omi Kumagai, Tomonori Kume, Runi anak Sylvester Pungga, Tohru Nakashizuka, Takao Itioka, Shoko Sakai

## Abstract

Supra-annually synchronized flowering events occurring in tropical forests in Southeast Asia, known as general flowering (GF), are “spectacular and mysterious” forest events. Recently, studies that combined novel molecular techniques and model-based theoretical approaches suggested that cool temperature and drought synergistically drove GF. Although the novel approaches advanced our understanding of GF, it is usually difficult to know whether the mathematical formulation reasonably well represents the complex and dynamic processes involved in GF. In the present study, we collected a 17-year set of community-wide phenology data during 1993 to 2011 from Lambir Hills National Park in Borneo, Malaysia, and analyzed it using a model-free approach, empirical dynamic modeling (EDM), that does not rely on specific assumptions about the underlying mechanisms, to overcome and complement the previous limitations. By using EDM, we found that GF in the forest in Lambir Hills National Park is synergistically driven by cool air temperature and drought, which is consistent with the previous studies. Also, we found that cumulative meteorological variables, rather than instantaneous values, drive GF with delayed effects, which is also consistent with the previous studies. Interestingly, the present study showed that effects of cumulative meteorological variables on GF changed through time, which implies that the relationship between GF and meteorological variables may be influenced by other factors such as plant/soil nutrient resource dynamics. Future studies integrating novel mathematical/statistical frameworks, long-term and large spatial scale ecosystem monitoring and molecular phenology data are promising for better understanding and forecasting of GF events in tropical forests in Southeast Asia.

## Introduction

Supra-annually synchronized flowering events occurring in tropical forests in Southeast Asia, known as general flowering (GF) or community-level masting, are “spectacular and mysterious” forest events (Ashton et al. 1988, Sakai 2002, Sakai et al. 2006). GF is a masting phenomenon unique to Asian dipterocarp forests, and it is unique in involving the synchronization of flowering/fruiting across diverse plant groups (Ashton et al. 1988, Sakai 2002). Plants including most dipterocarps and many other plant groups flower over roughly a 3-month period, and produce a large number of flowers (and subsequently, fruits) during the event. Conversely, flowers are rare between GF events especially in the canopy layer, while many understory herbs and treelets may flower more frequently. Therefore, GF plays an important role in plant reproduction and succession. In addition to its influence on plant communities, GF also influences forest animal communities, including pollinators and seed predators, through trophic cascades because plants provide a large amount of nutrient resources to forest animals (e.g. Curran and Leighton 2000, Itioka et al. 2001, Te Wong et al. 2005, Nakagawa et al. 2007, Iku et al. 2017). As a result, GF influences overall dynamics of forest ecosystems, and thus understanding GF mechanisms and predicting GF events are important for tropical forest conservation and management.

Because GF occurs over a large spatial scale and across a large number of plant groups, it is reasonable to assume that major drivers of GF are environmental rather than local biotic interactions. However, environmental cues that triggered GF have not been fully understood despite intense efforts of theoretical and empirical ecologists. Researchers have traditionally investigated potential environmental cues using observation and simple correlation analysis. For example, using simple correlations between environmental variables and the proportion of flowering and fruiting tree individuals, Sakai (2006) and Brearley (2007) independently suggested that irregular drought caused by El Niño southern oscillation (ENSO) events trigger GF. Other researchers suggested that low air temperature and nutrient resources (e.g., phosphorus) are important cues (Ashton et al. 1988, Ichie 2013), but detailed investigations of how these factors influence GF are still ongoing.

Recently, several research groups addressed this question by using advanced molecular techniques and theoretical approaches. For example, Kobayashi et al. (2013) investigated gene expression patterns of buds of *Shorea beccariana* before and during the flowering event in Lambir Hills National Park in Borneo, Malaysia. They found that expression levels of flowering-related genes, drought-responsive genes and sucrose-induced genes showed significant changes before flowering. Chen et al. (2017) analyzed 13 years of weekly flowering records collected in Pasoh Research Forest in Peninsular Malaysia using a mathematical modeling approach, and showed that drought and cool temperature synergistically best explain the timing of flowering events of *Shorea* species. Furthermore, Yeoh et al. (2017) used molecular phenology data collected from two *Shorea* species in Pasoh Research Forest to answer the question. The mathematical modeling of the molecular phenology data suggested that cool air temperature and drought synergistically influence the gene expression pattern of the *Shorea* (Yeoh et al. 2017). Molecular phenology, also known as the “ecological transcriptome” is the analyses of plant transcriptome time series collected from natural ecosystems, and has recently been recognized to be a powerful tool to understand causes and molecular mechanisms of plant phenology in nature (Aikawa et al. 2010, Kudoh 2016, Yeoh et al. 2017). These recent studies that utilized novel techniques suggested that several factors, rather than a single factor, are synergistically act as triggers of GF in Southeast Asia.

Although these advanced and integrated approaches have helped to advance our understanding of GF events, there still are several important limitations. First, although mathematical modeling that explicitly constructed model structures reasonably well explained GF events and/or gene expression patterns in the previous studies, it is difficult to know whether the mathematical formulations (i.e., model equations) reasonably well represented the complex and dynamic processes involved in GF. In other words, misspecifications of the model structure may lead us to wrong conclusions, but unfortunately, specifying model structures correctly is often difficult for complex, natural ecological dynamics. Second, in terms of gene expression data, measuring gene expression patterns from tropical tree species is still time-, labor-and money-consuming work, and thus, the molecular phenology data can often be collected from only a small number of individuals. Considering that GF is a community-wide phenomenon, analyzing community-wide data would provide important insights and reveal different aspects of GF mechanisms.

In the present study, we collected 17 years of community-wide phenology data from Lambir Hills National Park in Borneo, Malaysia, and analyzed it using a model-free approach to overcome and complement the limitations of earlier approaches. The long-term phenology data was collected by direct visual census (Sakai et al. 2006); the monitored plant individuals were first identified and the number of flowering plant individuals was counted fortnightly during June 1993 to January 2011 (i.e., the index of GF was “the proportion of flowering plant individuals in the forest”). Then, the time series data was analyzed using empirical dynamic modeling (EDM) (Sugihara et al. 2012, Ye et al. 2015a, Deyle et al. 2016). Instead of assuming a set of equations that govern the dynamics, EDM recovers system dynamics from the time series trajectory (for a brief introduction, see Chang et al. 2017 and Methods section). This unique feature enables us to analyze time series without assuming a specific model structure, and thus is often suitable for analyzing and predicting dynamics of complex systems such as natural ecosystems. In the time series analysis, we particularly focused on the influences of air temperature and rainfall because these two factors have been suggested to be dominant factors that trigger GF in many studies (Ashton et al. 1988, Sakai et al. 2006, Brearley et al. 2007, Kobayashi et al. 2013, Chen et al. 2017, Yeoh et al. 2017). The community-wide and model-free analysis, which we present here, was lacking in the previous studies, and we tested whether the “cool air temperature and drought hypothesis” was validated using our approach.

## Methods

### Study site

The study site is in a lowland dipterocarp forest at Lambir Hills National Park, Borneo (4° 20′ N, 113°50′ E, 150–250 m a.s.l.) (Roubik et al. 2005). Rainfall data were collected at the Bukit Lambir Station of the Department of Irrigation and Drainage (DID), Sarawak, Malaysia, within the park approximately 3 km northwest of the study site. From 1985–2003, the average annual rainfall was *ca*. 2700 mm, and the annual rainfall ranged from 2000 to 3800 mm. Seasonal variation was significant but low, and the area occasionally had droughts with biological consequences (Harrison 2000, Nakagawa et al. 2000, 2019, Itioka and Yamauti 2004, Kishimoto-Yamada et al. 2009).

### Time series of general flowering

Plants were observed using a canopy observation system (tree towers and walkways) constructed at the center of the Canopy Biology Plot (8 ha, 200 × 400 m) (Roubik et al. 2005). The Canopy Biology Plot includes humult and udult soils (sandy clay, light clay, or heavy clay in texture), several ridges and valleys, and closed (mature-stage) forests and canopy gaps. In each census, presence or absence of flowers on focal plants was recorded fortnightly (Sakai et al. 2006). After excluding plants with missing values, records of 204 individuals from June 6^th^ 1993 to January 5^th^ 2011 were used in the time series. The monitored plant individuals belong to 38 families, 80 genera and 133 species, and dominant families are as follows: Dipterocarpaceae (*N*=74; genera *Shorea*, *Dipterocarpus*, *Dryobalanops* and *Vatica* were included), Euphorbiaceae (*N*=23), Leguminosae (*N*=17), Annonaceae (*N*=8), Burseraceae (*N*=8), Myristicaceae (*N*=7) and other families (*N*=67). General flowering events were observed seven times during the census period (Fig. 1a). During the 17 years and 7 months census period, the total number of time points used for the general flowering time series was 423 (i.e., fortnightly census for 211 months).

**Figure 1:**
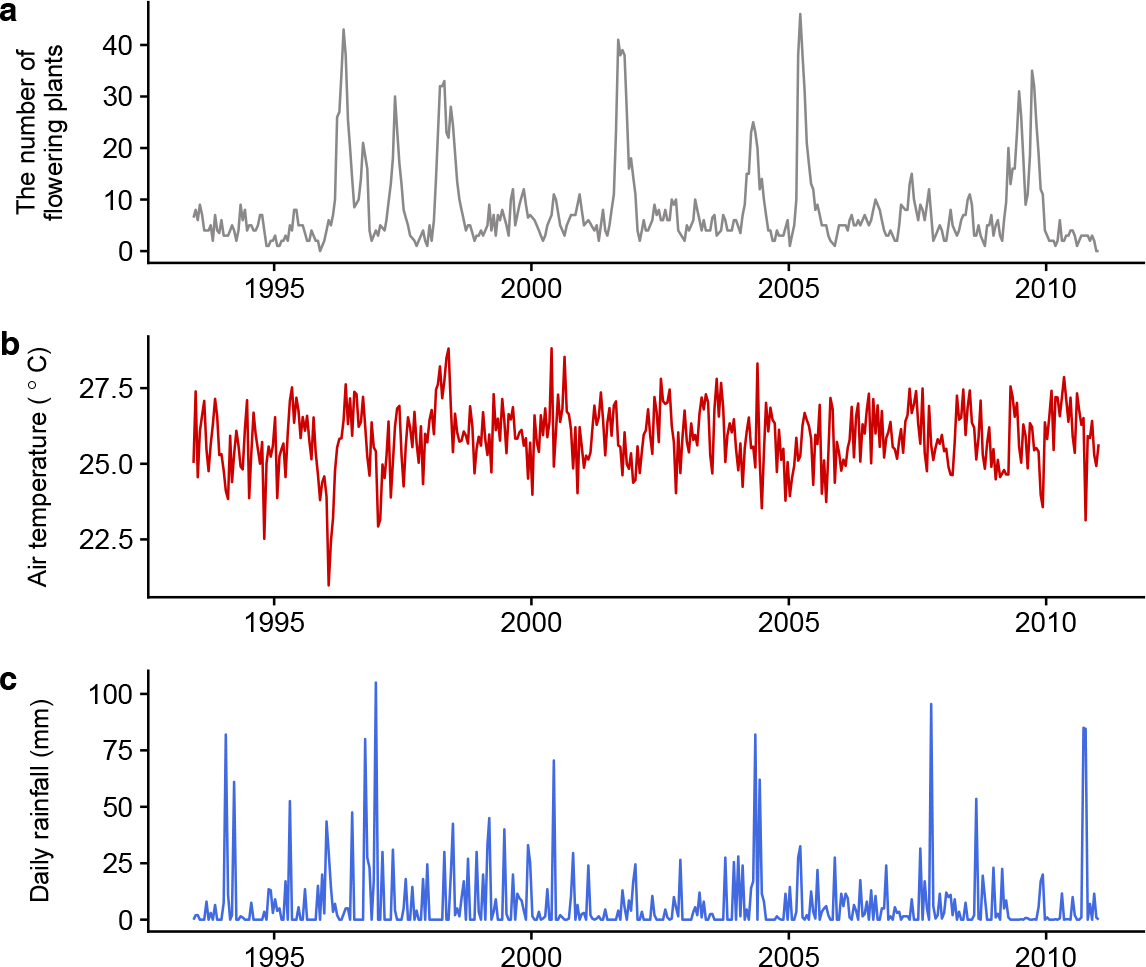
Time series of general flowring and climate variables. (**a**) The number of flowering plant individuals. The total number of monitored plant individuals was 204. (**b**) Daily mean air temperature (^◦^C). (**c**) Daily rainfall (mm).

### Meteorological time series

From 1993 to 1998, temperature was monitored every 30 min using a temperature/humidity sensor (E7050– 10, Yakogawa Weathac Corp., Tokyo, Japan) on a tower 35 m above the ground located in the Canopy Biology Plot (Sakai et al. 2006). From 2000 to 2011, temperature was monitored every 10 min using a thermohygrograph (HMP35A, Visala Co., Helsinki, Finland) installed at the height of 76 m on a crane tower located approximately 0.5 km from the plot (Kume et al. 2011). For rainfall from 1993 to 1999, data collected daily at the Bukit Lambir Station of the Department of Irrigation and Drainage (DID), Sarawak, Malaysia, within the park approximately 3 km northwest of the study site, was used. From 2000 to 2011, rainfall was monitored using a tipping bucket rain gauge (RS102, Ogasawara Keiki, Tokyo, Japan) at the top of the crane, 85.8 m above the ground (Kume et al. 2011).

In the analysis, we used the observed meteorological data when it was available. When observed data was not available, the data was complemented by a reanalyzed and gridded four-dimensional meteorology dataset, JRA-55 (Kobayashi et al. 2015) for the nearest grid point 3° 45′N, 113° 45′E. In total, air temperature data at 62 time points (with two-week intervals) was not available among the total 423 time points and was complemented by using the simulated air temperature data (a major missing period for air temperature data was from October 1998 to January 2001; 31 time points). For rainfall data, there was no missing data and thus only observed data was used for the analyses. Although the study site is an aseasonal tropical rain forest, there are variations in daily mean air temperature and rainfall (Fig. 1b, c; Kume et al. 2011).

### Calculations of cumulative air temperature and rainfall

Meteorological variables were first converted to daily values (i.e., daily mean air temperature and daily rainfall) during the census term (i.e., from June 1993 to January 2011), and the cumulative values for the daily variables were calculated. To examine the most influential cumulative duration, we calculated the cumulative values from 7-days to 364-days with 7-day intervals. Thus, we had 52 cumulative values × two meteorological variables = 104 variables to be examined by empirical dynamic modeling (see the section below).

### Framework of empirical dynamic modeling (EDM): State space reconstruction (SSR)

Time series can be defined as any set of sequential observations of the system state, and the dynamic behaviors can be delineated as a trajectory of a state over time in a multidimensional state space by plotting time series. Time series taken from ecosystems (i.e., ecological time series) can be used to trace out trajectories of the system, which provide a large amount of information on ecosystem dynamics. For example, if one has performed sequential observations on a three-species ecological system, e.g., grasses (primary producer), rabbits (consumer) and foxes (predator), then the dynamics of the three-species system can be reconstructed by plotting time series of grasses, rabbits, and foxes along the *x*, *y*, and *z* axis, respectively, in a three-dimensional state space. The motion of the three-dimensional vectors can be understood as the system behavior.

In a natural ecosystem, however, it is almost impossible to collect time series of all potentially important variables involved in a target system. Fortunately and counterintuitively, Takens (1981) offered a theoretical basis to solve this problem: a mathematical theorem, Takens’ embedding theorem, demonstrated that a shadow version of the attractor (motion of vectors in a state space) can be reconstructed by a single observed time series (for example, the time series of grasses, *x*). In other words, delineation of trajectories, originally constructed using multivariables, can be possible even if a time series is available only for a single variable (Takens 1981, Sauer et al. 1991). To embed such a single time series (with an equal sampling interval), vectors in the putative phase space are formed from time-delayed values of the time series, {*x* (*t*), *x* (*t − τ*), *x* (*t* − 2*τ*), …, *x* (*t* − [*E* − 1] *τ*)}, where *E* is the embedding dimension, and *τ* is the time lag. This procedure, the reconstruction of the original dynamics, is known as State Space Reconstruction (SSR) (Takens 1981, Deyle and Sugihara 2011).

Recently developed tools for nonlinear time series analysis, which are specifically designed to analyze state-dependent behavior of dynamical systems, called Empirical Dynamic Modeling (EDM), are rooted in SSR (Sugihara and May 1990, Sugihara 1994, Dixon et al. 1999, Ye et al. 2015a, 2015b, Deyle et al. 2016, Ye and Sugihara 2016, Kitayama et al. 2018, Ushio et al. 2018). These methods do not assume any set of equations governing the system, and thus are suitable for analyzing complex systems, for which it is often difficult to make reasonable *a priori* assumptions about their underlying mechanisms. Instead of assuming a set of specific equations, EDM recovers the dynamics (and potentially, underlying mechanism) directly from time series data, and is thus particularly useful for forecasting ecological time series, which are otherwise often difficult to forecast.

### Convergent cross mapping between general flowering and meteorological variables

In order to detect causation between GF and meteorological variables, we used convergent cross mapping (CCM) (Sugihara et al. 2012). An important consequence of the SSR theorems is that if two variables are part of the same dynamical system, then the reconstructed state spaces of the two variables will represent topologically the same attractor (with a one-to-one mapping between reconstructed attractors). Therefore, it is possible to predict the current state of a variable using time lags of another variable. We can look for the signature of a causal variable in the time series of an effect variable by testing whether there is a correspondence between their reconstructed state spaces (i.e., cross mapping). This cross map technique can be used to detect causation between variables. Cross-map skill can be evaluated by a correlation coefficient (*ρ*), or mean absolute error (MAE) or root mean square error (RMSE) between observed and predicted values by cross mapping. In addition, the cross-map skill will increase as the library length (i.e., the number of points in the reconstructed state space of an effect variable) increases if two variables are causally related (i.e., convergence), and the convergence is a practical criterion of causality, and thus the analysis is called convergent cross mapping (CCM).

In the present study, cross mapping from one variable to another was performed using simplex projection. How many time lags are taken in SSR (i.e, best embedding dimension; *E*) is determined by simplex projection using RMSE as an index of forecasting skill. Throughout the analysis, *E* = 6 was used for GF time series. More detailed algorithms about simplex projection and cross mapping can be found in previous studies (Sugihara and May 1990, Sugihara et al. 2012, Chang et al. 2017).

In CCM analysis, we also considered the time lag between GF time series and potential causal time series (i.e., cumulative meteorological variables). This can be done by using “lagged CCM” (Ye et al. 2015b). For normal CCM, correspondence between reconstructed state space (i.e., cross-mapping) is checked using the same time point. In other words, information embedded in an effect time series at time *t* may be used to predict the state of a potential causal time series at time *t*. This idea can easily be extended to examine time-delayed influence between time series by asking the following question: is it possible to predict the state of a potential causal time series at time *t − tp* (*tp* is a time delay) by using information embedded in an effect time series at time *t*? Ye et al. (2015b) showed that lagged CCM is effective to determine the effect time delay between variables. In the present study, we examined the effect time delay from 0-days to 336-days with 14-day intervals.

In this present study, the significance of CCM is judged by comparing convergence in the cross-map skill of Fourier surrogates and original time series. More specifically, first, 1,000 surrogate time series for one original time series are generated. Second, the convergence of the cross-map skill (i.e., the difference between cross-map skills (RMSE) at the minimum and maximum library lengths, denoted by ∆RMSE) is calculated for 1,000 surrogate time series and the original time series. If the number of surrogates that shows a higher convergence is less than 50 (i.e., 5% of the surrogates), the cross mapping is judged as significant.

### Multivariate S-map method

The multivariate S-map (sequential locally weighted global linear map) method allows quantifications of dynamic (i.e., time-varying) interactions (Sugihara 1994, Deyle et al. 2016). Consider a system that has *E* different interacting variables, and assume that the state space at time *t* is given by *x*(*t*) = {*x_1_*(*t*), *x_2_*(*t*), …, *x_E_*(*t*)}. For each target time point *t*\*, the S-map method produces a local linear model ***C*** that predicts the future value *x_1_*(*t*\*+*p*) from the multivariate reconstructed state space vector *x*(*t*\*). That is,

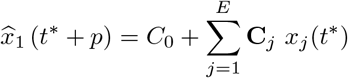

where 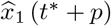 is a predicted value of *x*_1_ at time *t*^*^+*p*, and *C*_0_ is an intercept of the linear model. The linear model is fit to the other vectors in the state space. However, points that are close to the target point, *x*(*t^*^*), are given greater weighting. The model ***C*** is the singular value decomposition (SVD) solution to the equation

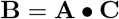

where ***B*** is an *n*-dimensional vector (*n* is the number of observations) of the weighted future values of *x_1_*(*t_i_*) for each historical point, *t_i_*, given by

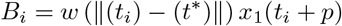

and ***A*** is then *n × E* dimensional matrix given by

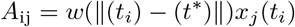

The weighting function *w* is defined by

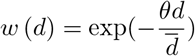

which is tuned by the nonlinear parameter *θ* ≧ 0 and normalised by the average distance between *x*(*t*\*) and the other historical points,

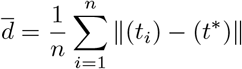

||*x* – *y*|| is the Euclidian distance between two vectors in the *E*-dimensional state space. Note that the model ***C*** is separately calculated (and thus potentially unique) for each time point, *t*. As recently shown, ***C***, the coefficients of the local linear model, are a proxy for the interaction strength between variables (Deyle et al. 2016).

### Computation and result visualization

Simplex projection, S-map and CCM were performed using “rEDM” package (version 0.7.1) (Ye et al. 2015a, 2018), results were visualized using “ggplot2” (Wickham 2009), “cowplot” (Wilke 2017), and “ggsci” (Xiao 2018) packages. All statistical analyses were performed in the free statistical environment R3.4.3 (R Core Team 2017).

### Code and data availability

Our time series data and computing codes include unpublished works that involved multiple researchers and institutions. Therefore, the data and codes are currently not deposited in a public archive such as Dryad, GitHub or Zenodo. They will eventually be available as a Data paper and/or original research papers. At present, all data and code are available upon reasonable request.

## Results and Discussion

### Detection of causal factors of general flowering

According to the lagged CCM, 7-days cumulative air temperature with 28-days delay and 7-days cumulative rainfall with 42-days delay most strongly cause the number of flowering plant individuals in the Lambir forest plot (Fig. 2, *P* < 0.05). In the analysis, we relied on ∆RMSE, but qualitatively similar results were obtained even when using ∆*ρ* and ∆MAE (Fig. S1–2). Furthermore, including 7-days cumulative air temperature with 28-days delay and 7-days cumulative rainfall with 42-days delay in the multivariate S-map improved the forecasting skill (RMSE and *ρ*) of GF compared with the univariate model and models with air temperature only or rainfall only (Fig. 3). These results suggest that air temperature and rainfall synergistically, not independently, influence GF in the study site.

**Figure 2:**
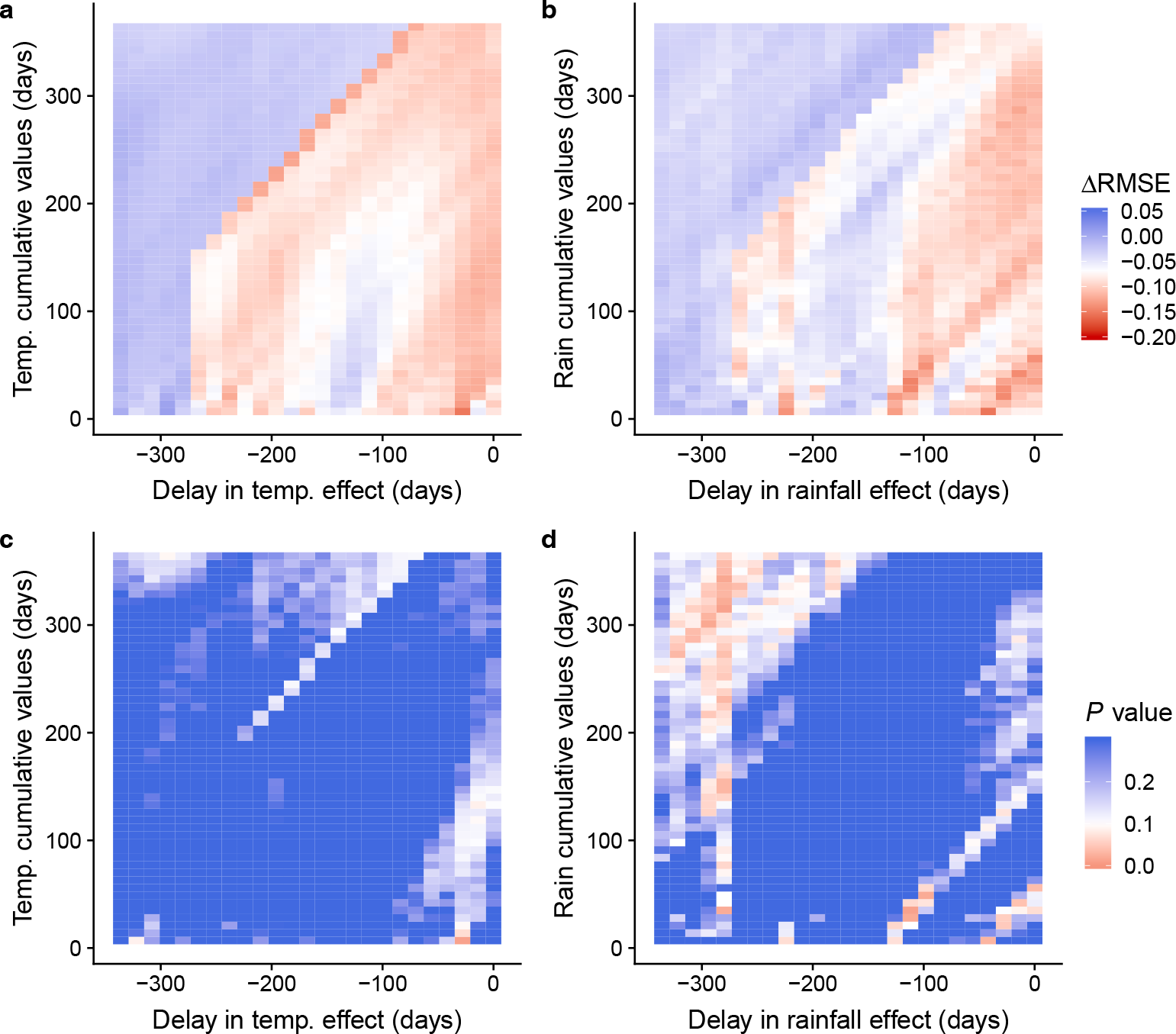
Results of convergent cross mapping (CCM). Improvement of forecasting skill (measured by RMSE [Root Mean Square Error]) by CCM for different combinations of effect time-delay and cumulative values of air temperature (**a**) and rainfall (**b**). *P* values for different combinations of effect time-delay and cumulative values of air temperature (**c**) and rainfall (**d**).

**Figure 3:**
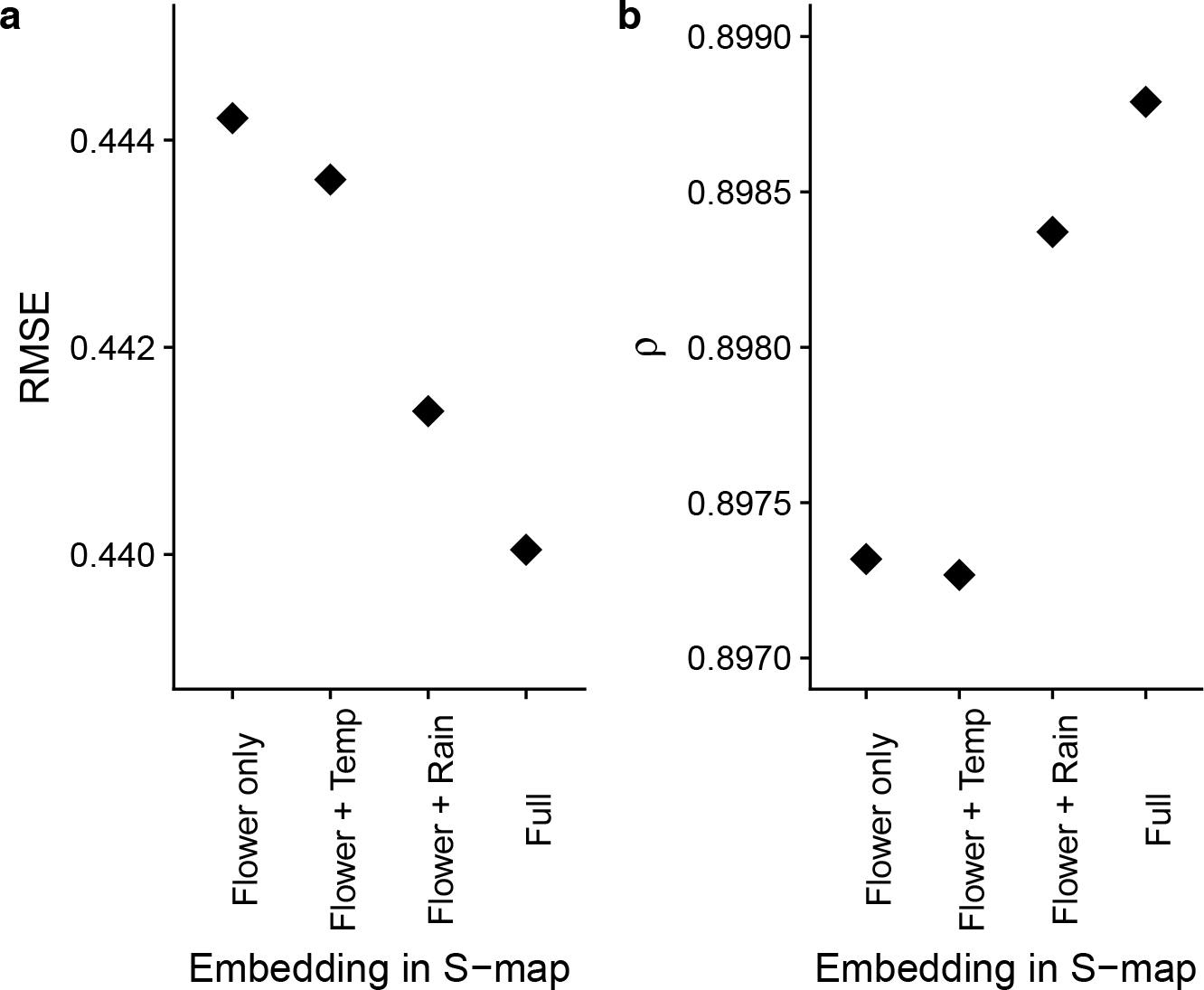
Improvements of forecasting skill by the univariate/multivariate S-map method. (**a**) Root mean square error (RMSE) by the S-map methods that used four different embeddings. “Flower only” indicates that only GF index was used to forecast GF index (i.e., embedding = {GF_t_, …, GF_t − 5_}). “Flower + Temp” and “Flower + Rain” indicates embedding was {GF_t_, …, GF_t – 4_, Cum.T emp} and {GF_t_, …, GF_t – 4_, Cum.Rainf all}, respectively. “Full” indicates that the embedding was {GF_t_, …, GF_t – 3_, Cum.T emp, Cum.Rainf all} (**b**) Correlation coefficient (ρ) by the S-map methods that used four different embeddings.

In addition to 7-days cumulative air temperature with 28-days delay and 7-days cumulative rainfall with 42-days delays, other cumulative values with different delays may have causal influences on the number of flowering plant individuals in the forest (Fig. 2). This suggests that detailed mechanisms of the influences of air temperature and rainfall may include several different pathways, e.g., air temperature in one day may consequently influence the number of flowering plant individuals in different days. However, including all potential cumulative and delay values makes a model extremely complex and reduces the model interpretability. Therefore, in the present study, we focus on the influences of 7-days cumulative air temperature with 28-days delay and 7-days cumulative rainfall with 42-days delays as dominant environmental drivers of GF in the subsequent analysis.

### Influences of air temperature and rainfall

Influences of 7-days cumulative air temperature and rainfall were quantified using the multivariate S-map method. In general, 7-days cumulative air temperature and rainfall have negative influences on GF (Fig. 4a, red and blue solid lines), indicating that decreased air temperature and drought increase the number of flowering plant individuals in the forest, which is consistent with the previous studies (Sakai et al. 2006, Kobayashi et al. 2013, Chen et al. 2017, Yeoh et al. 2017). However, though in general air temperature and rainfall have negative influences on GF, air temperature sometimes has positive influences on GF, i.e., increased air temperature increases the number of flowering plant individuals in the forest (Fig. 4a; the red line sometimes exceeds the zero value indicated by the dashed line).

**Figure 4:**
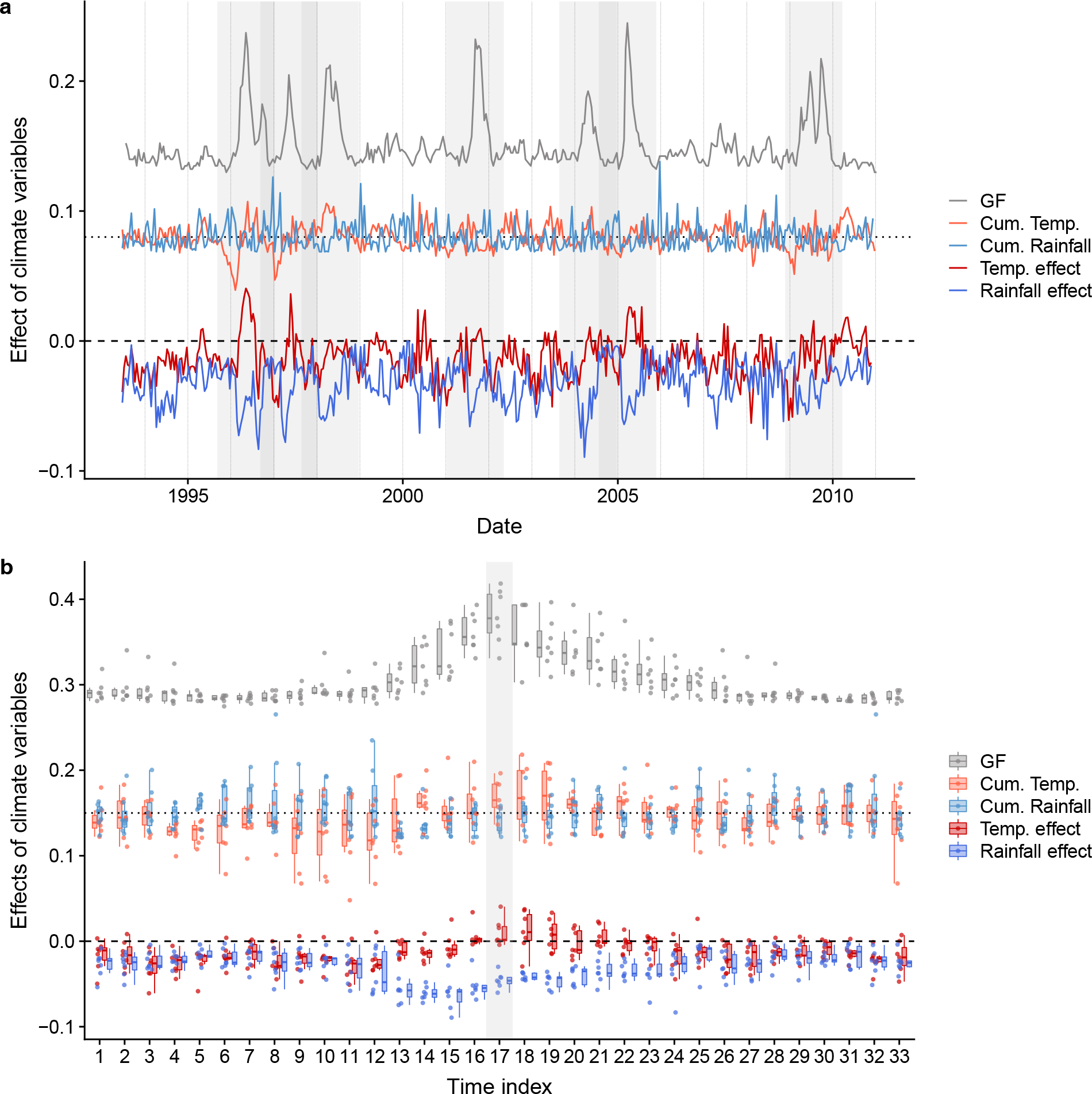
Time series of GF, cumulative climate variables and their influence on GF. (**a**) Standardized GF and cumulative temperature and rainfall were plotted. Values on the y-axis are only for effects of climate variables. Dotted and dashed horizontal lines indicate mean value for cumulative climate variables and zero for effects of climate variables, respectively. Different line colors indicate different time series categories. Vertical dotted lines correspond to one-year. Vertical shaded regions indicate the GF periods and were used to make panel **b**. (**b**) Time-windows of 8-montsh before and 6-months after each GF peak were extracted from panel **a** and plotted using box and jitter plots. Seven GF peroids, i.e., 1995/9-1997/1, 1996/9-1998/1, 1997/8-1998/12, 2001/1-2005/8, 2003/8-2004/12, 2004/7-2005/12, and 2008/11-2010/3, were extracted. Shaded region indicates the GF peak (time index 17). One time index corresponds to 2-weeks. Dotted and dashed horizontal lines indicate mean value for cumulative climate variables and zero for effects of climate variables, respectively. Note that effects of cumulative temperature and rainfall are delayed by 28-days and 42-days, respectively. For example, cumulative rainfall (and its effect) at time index 4 influence GF at time index 6.

In order to examine the detailed patterns of the influences of air temperature and rainfall on GF, a 14-month time window around each GF event (i.e., 8 months before GF and 6 months after GF) was extracted (Fig. 4b; time windows around seven GF periods were used, i.e., the periods of 1995/9–1997/1, 1996/9–1998/1, 1997/8–1998/12, 2001/1–2005/8, 2003/8–2004/12, 2004/7–2005/12 and 2008/11–2010/3, indicated by gray shading in Fig. 4a). According to the visualization, GF, air temperature and rainfall and their influences on GF showed consistent patterns across the seven GF events (Fig. 4b). The GF event generally started 2.5–3 months (i.e., time indices 12–13) before its peak (at time index 17 in Fig. 4b), and deceased air temperature was found during *ca*. 4 months before the initiation of GF (time indices 4–12 in Fig. 4b, orange points and boxplots). Therefore, a first cue of a GF event might be decreased air temperature rather than drought. Interestingly, increases in rainfall that coincided with decreases in air temperature were found *ca*. 4 months before the initiation of GF (time indices 5–6 in Fig. 4b, skyblue points and boxplots), which might be a previously overlooked sign of GF initiation. Then, in the GF increasing period, drought positively and strongly influenced the number of flowering plant individuals (time indices 12–16 in Fig. 4b, skyblue and blue points and boxplots). After the GF peak, slightly increased air temperature contributed to the maintenance of GF (time indices 17–20 in Fig. 4b, orange and red points and boxplots). Interestingly and importantly, these patterns were generally consistent across the seven GF events, and thus monitoring air temperature and rainfall may help to forecast GF.

In order to search for more detailed patterns, we examined the relationships between GF, air temperature, rainfall and their influences using scatter plots (Fig. 5). According to the analysis, there may be a threshold of cumulative rainfall to induce GF. GF happens only when 7-day cumulative rainfall values are below *ca*. 100 mm (Fig. 5a). This suggests that drought may be a necessary condition, but not a sufficient condition, for GF. On the other hand, there is no clear and linear relationship between GF and air temperature (Fig. 5b), suggesting that air temperature is, though significant, a less important factor of GF than drought. In addition, during GF periods, the influence of drought is stronger than that during other periods, and there are linear relationships between climate variables and their influences on GF (Fig. 5c, d). This suggests that the effects of climate factors on GF are not constant, but can change depending of the states of a forest ecosystem. Also, more severe drought, and/or lower air temperature have stronger influences on GF, and thus, they may induce a larger GF event.

**Figure 5:**
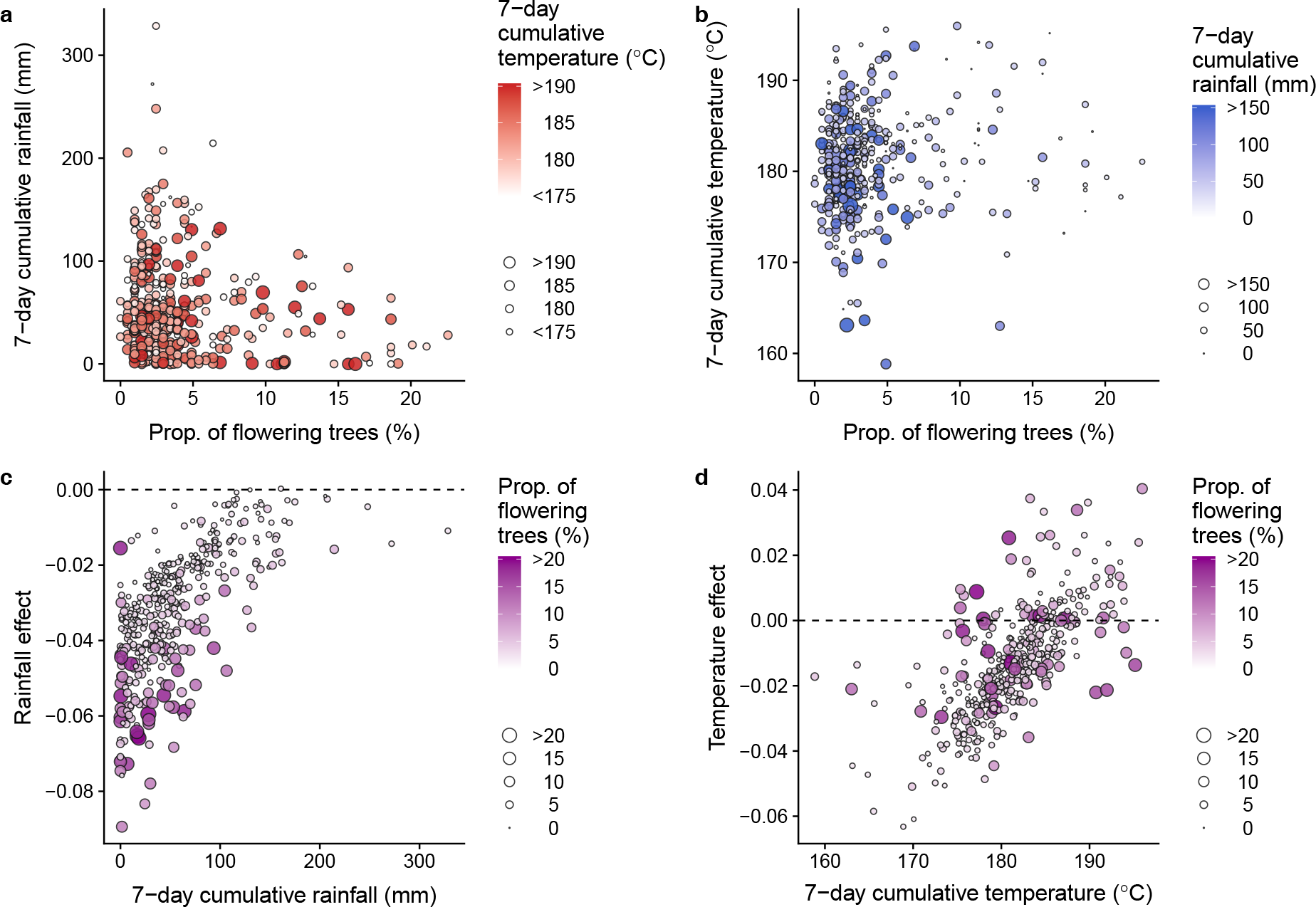
The relationships between the proportion of flowering plants, cumulative climate variables and effects of cumulative climate variables. (**a**) The relationship between the proportion of flowering plants and 7-day cumulative rainfall. Color density and point size indicate 7-day cumulative temperature. (**b**) The relationship between the proportion of flowering plants and 7-day cumulative temperature. Color density and point size indicate 7-day cumulative rainfall. (**c**) The relationship between 7-day cumulative rainfall and effects of cumulative rainfall. Color density and point size indicate the proportion of flowering plants. (**d**) The relationship between 7-day cumulative temperature and effects of cumulative temperature. Color density and point size indicate the proportion of flowering plants.

### Comparisons with previous studies using different approaches

Using a 17-year, fortnightly direct visual observation of GF and EDM, we showed that air temperature and rainfall synergistically and dynamically drive GF at a community-level in the tropical forest in Lambir Hills National Park in Borneo, Malaysia. This finding is consistent with the recent studies that suggested synergistic influences of air temperature and rainfall (Chen et al. 2017, Yeoh et al. 2017). Responses of expression levels of drought-and/or air temperature-related genes may be an important individual-level basis for the initiation of GF, as found in previous molecular studies (Kobayashi et al. 2013), but elucidating how these molecular-level responses scale up to community-level responses will need further studies.

Previous studies that analyzed flowering records by mathematical modeling suggested that “signal accumulation” (equivalent to our “cumulative values”) ranged from 54 to 90 days, and “days for flower development” (equivalent to our “effect time delay”) ranged from 43 to 96 days (Chen et al. 2017), and these values were longer than our results that showed 7-days cumulative air temperature with a 28-days effect delay and 7-days cumulative rainfall with a 42-days effect delay are the most influential climate variables. These differences may be partly due to differences in modeling approaches. When using a model-based approach, estimated parameter values depend on the model structure. In other words, if the model structure changes, estimated parameter values (i.e., days for signal accumulations and days for flower development) may change. The differences may also be due to the characteristics of the time series analyzed. In Chen (2017), species-specific flowering records were analyzed, but our time series are community-wide flowering records. Therefore, estimated cumulative days and effect delays were community-wide (or community-averaged) ones, and thus they may be different from species-specific parameters. Lastly, although our analysis focused on the most influential cumulative values and effect delays, longer cumulative values and effect delays are also suggested to be factors that influence GF (Fig. 2a, b), and the previous studies might have detected such longer values as the most influential values. Altogether, it is not surprising that we found several differences in the estimated parameter values, considering the differences in modeling approaches and time series characteristics.

One of the interesting findings of our study is that effects of air temperature and rainfall may change depending on the states of the forest ecosystem (Figs. 4, 5), which was not pointed out in the previous studies. The time-varying effects of climate may be reasonable because factors other than climate (e.g., tree physiological conditions such as the level of internal nutrient resources) are suggested to be an important factor to determine the flowering timing (Ichie 2013). For example, when sufficient phosphorus and/or nitrogen is not stored in the tree body, climate cues, e.g., drought and low air temperature, would not induce flowering. Also, the best embedding dimension of GF time series was estimated to be 6 (*E* = 6), suggesting that the number of potential factors that influence GF might be more than 2 (i.e., air temperature and rainfall) (Takens 1981). Incorporating other potential factors (e.g., plant and soil resource dynamics, tree root-fungal associations and other meteorological variables) might further improve the skill of forecasting GF, which would contribute to better understanding of GF mechanisms.

### Conclusions

Using a long-term phenology monitoring data set and empirical dynamic modeling, we showed that GF in the forest in Lambir Hills National Park is synergistically driven by decreased air temperature and drought. These findings are consistent with those of previous monitoring, molecular and statistical studies (Sakai et al. 2006, Kobayashi et al. 2013, Chen et al. 2017, Yeoh et al. 2017). Several studies, including the present study, suggested that cumulative meteorological variables, rather than instantaneous values, with some effects of delays may drive GF, but robust estimations of the values of parameters for the accumulation and delay effects will need further studies. Unlike previous studies, the present study showed for the first time that the effect of meteorological variables on GF may change over time, which implies that some other factors such as plant/soil nutrient resource dynamics may be involved in GF. Integrating novel mathematical/statistical frameworks, long-term and large spatial scale ecosystem monitoring and large-scale molecular phenology is promising for better understanding and forecasting GF events in tropical forests in Southeast Asia.

## Acknowledgements

This study was conducted in accordance with memorandums of understanding signed in 2005 by the Sarawak Forestry Corporation (SFC, Kuching, Malaysia) and the Japan Research Consortium for Tropical Forests in Sarawak (JRCTS, Sendai, Japan), and in 2012 by the Sarawak Forest Department (SFD, Kuching, Malaysia) and JRCTS. We thank Mohd Shahbudin Sabki and other staff of SFD, Lucy Chong and other staff of SFC, and staff of Lambir National Park for their support for our study, and Chih-hao Hsieh for discussion about EDM. This study was financially supported by Grants-in-Aid (No. 16H04830 to S.S.) from the Japanese Ministry of Education, Science and Culture.

