## Supporting information for "Dynamic and synergistic influences of air temperature and rainfall on general flowering in a Bornean lowland tropical forest"

### Contents:

- **Figure S1**| Results of convergent cross mapping (CCM) evaluated by correlation coefficient, $\rho$
- **Figure S2**| Results of convergent cross mapping (CCM) evaluated by mean absolute error (MAE)

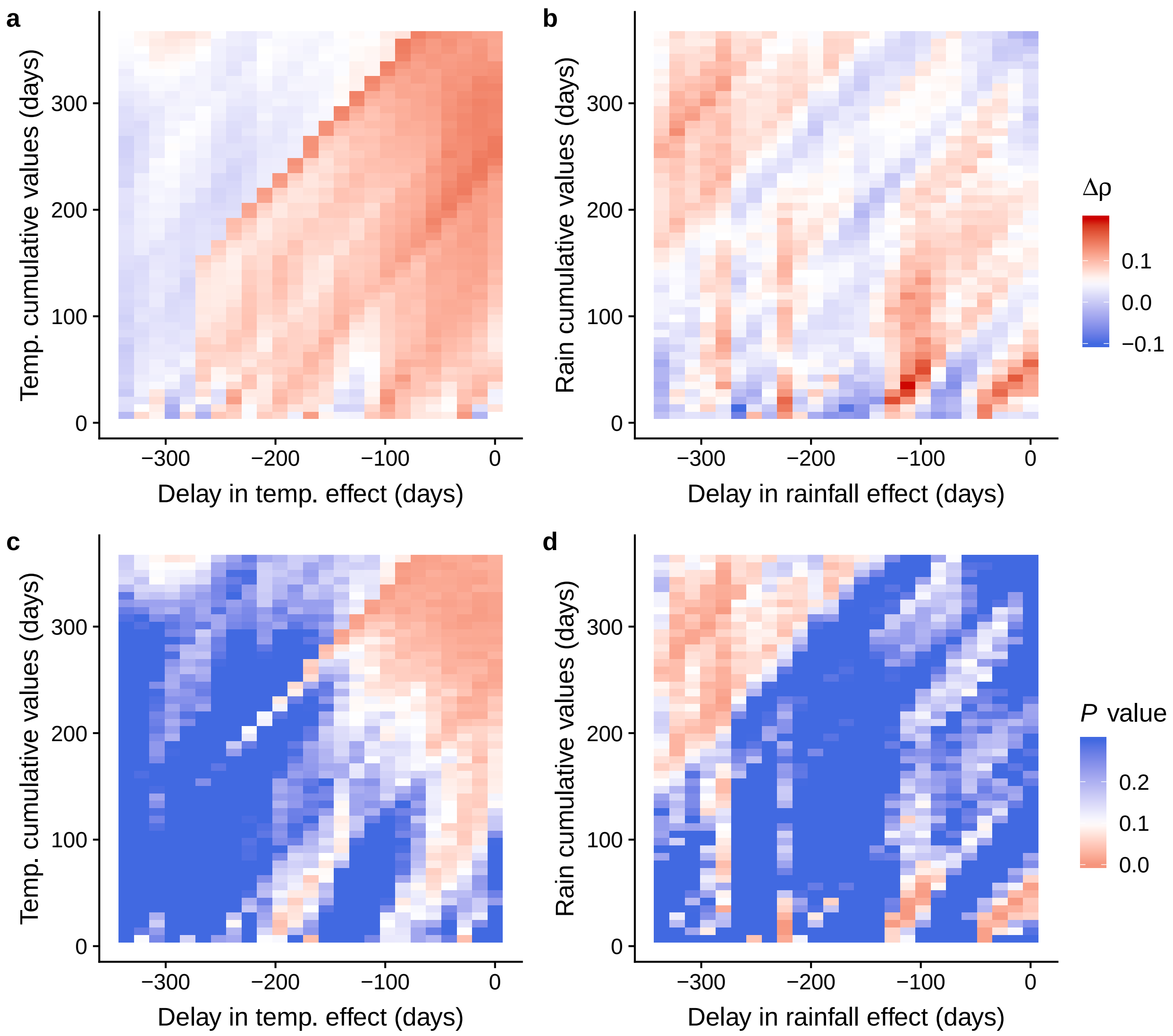


**Figure S1**| Results of convergent cross mapping (CCM). Improvement of forecasting skill (measured by correlation coefficient, $\rho$) by CCM for different combinations of effect time-delay and cumulative values of air temperature (**a**) and rainfall (**b**). $P$ values for different combinations of effect time-delay and cumulative values of air temperature (**c**) and rainfall (**d**).

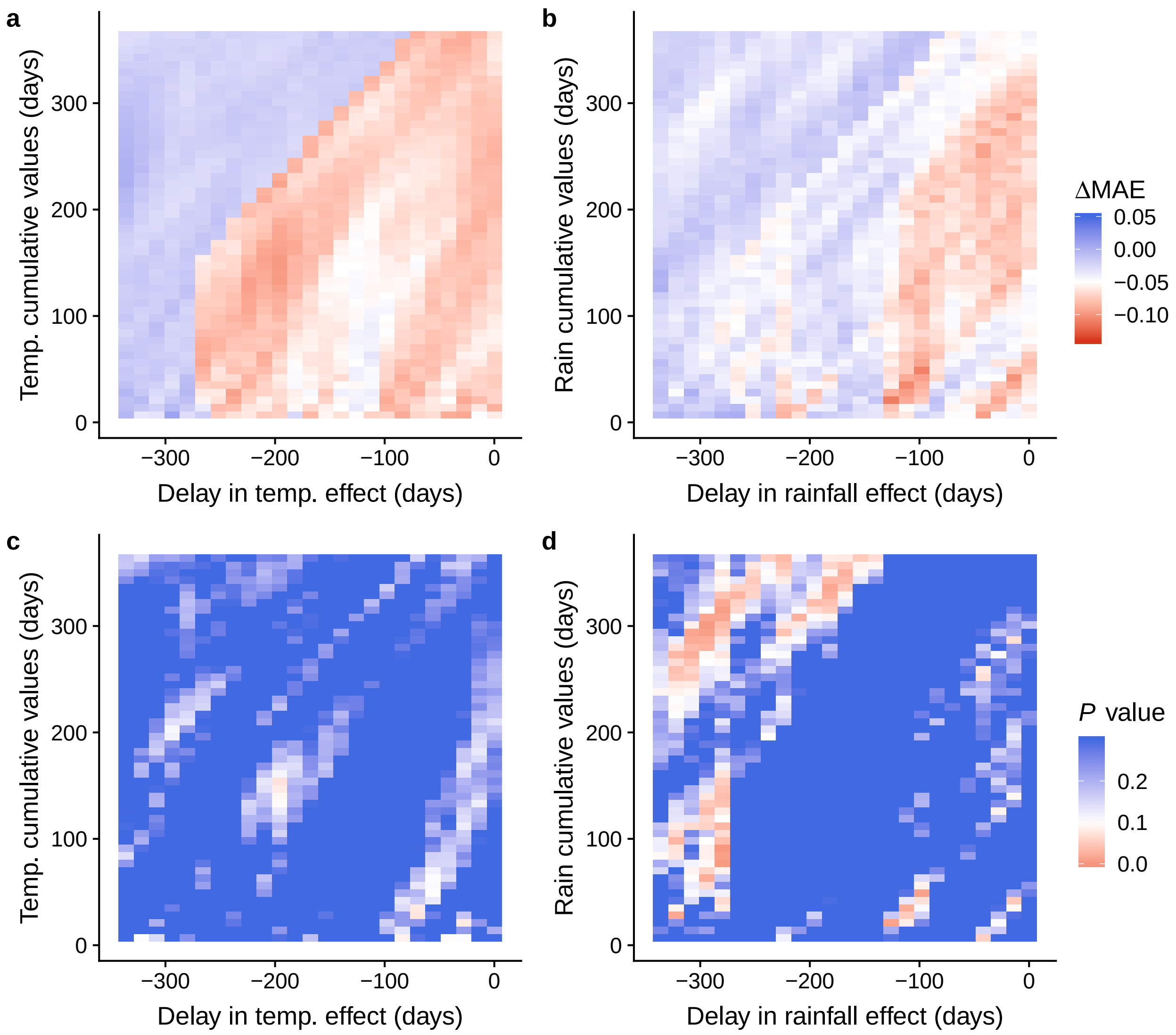


**Figure S2**| Results of convergent cross mapping (CCM). Improvement of forecasting skill (measured by correlation coefficient, MAE [Mean Absolute Error]) by CCM for different combinations of effect time-delay and cumulative values of air temperature (**a**) and rainfall (**b**). *P* values for different combinations of effect time-delay and cumulative values of air temperature (**c**) and rainfall (**d**).
